# Data-driven structural analysis of Small Cell Lung Cancer transcription factor network suggests potential subtype regulators and transition pathways

**DOI:** 10.1101/2023.04.01.535226

**Authors:** Mustafa Ozen, Carlos F. Lopez

## Abstract

Small Cell Lung Cancer (SCLC) is an aggressive disease and challenging to treat due to its mixture of transcriptional subtypes and subtype transitions. Transcription factor (TF) networks have been the focus of studies to identify SCLC subtype regulators via systems approaches. Yet, their structures, which can provide clues on subtype drivers and transitions, are barely investigated. Here, we analyze the structure of an SCLC TF network by using graph theory concepts and identify its structurally important components responsible for complex signal processing, called hubs. We show that the hubs of the network are regulators of different SCLC subtypes by analyzing first the unbiased network structure and then integrating RNA-seq data as weights assigned to each interaction. Data-driven analysis emphasizes MYC as a hub, consistent with recent reports. Furthermore, we hypothesize that the pathways connecting functionally distinct hubs may control subtype transitions and test this hypothesis via network simulations on a candidate pathway and observe subtype transition. Overall, structural analyses of complex networks can identify their functionally important components and pathways driving the network dynamics. Such analyses can be an initial step for generating hypotheses and can guide the discovery of target pathways whose perturbation may change the network dynamics phenotypically.

## INTRODUCTION

Throughout their evolution, cells differentiate and specialize into different subtypes, that are often controlled by underlying molecular-level mechanisms [1-3]. This process is generally pictured by the famous metaphor that is a ball rolling down a hill, called the Waddington Landscape [4]. Analogous to a ball rolling down a hill, which may change its direction by the effect of obstacles in its way, lose its kinetic energy, slow down, and eventually reside at a stable point, cells may change their trajectories and differentiate to different subtypes due to some regulatory or evolutional triggers while they are maturing. Similarly, due to abnormalities, stochasticity, or other unknown reasons, they may diverge from their trajectories and become cancerous cells [5]. Moreover, cancerous cells may also evolve and differentiate into other subtypes [6-8]. Therefore, developing effective treatments for cancer has been a challenge due to heterogeneous cell subpopulations that appear within a tumor. Genetic or non-genetic mechanisms can drive the cancerous cell subpopulations via plasticity, drug-induced selection, or state transitions between the subtypes and have them escape the treatment or recur with a resistance to the treatment [9-11], which is the case in multiple cancer types such as breast cancer [12,13], melanoma [14], and Small Cell Lung Cancer (SCLC) [15-20].

SCLC is an extremely aggressive disease with a low survival rate [21-25] (7% 5-year survival as of 2022 [26]). Although it was characterized as molecularly homogeneous due to loss of TP53 and RB1, and neuroendocrine/epithelial differentiation [27,28], SCLC was shown to be heterogeneous [29-37] by the identification of its mixtures of transcriptional subtypes such as *neuroendocrine* (NE) stem-cell-like subtype centered on the expression of the transcription factors ASCL1 and NEUROD1 [35] and *non-neuroendocrine* (NON-NE) subtype centered on the expression of the transcription factor YAP1 [36]. Overall, the SCLC subtypes have been classified into four classes SCLC-A (also labeled as NE), SCLC-N (also labeled as NEv1), SCLC-Y (also labeled as NON-NE), and SCLC-P defined by the expression of the transcription factors ASCL1 (A), NEUROD1 (N), YAP1(Y), and POU2F3 (P), respectively [29-37]. Recently, the fifth subtype has also been proposed named SCLC-A2 (also labeled as NEv2) which is driven by ASCL1 but distinct from the SCLC-A neuroendocrine subtype [38]. The disease seems to start by including the NE type, and then the cancerous cell population begins to include the NON-NE subtype, which is more treatment-resistant [34,39,40]. In addition to various subtypes with different levels of resistance to treatment, such transitions between the subtypes further complicate the treatment of the disease. Therefore, understanding molecular heterogeneity in SCLC is essential for developing more precise, tailored treatments to cure the pathology.

Transcription factor (TF) networks have been the focus of the studies to understand the mechanism of the disease and to identify different SCLC subtypes as they are associated with the overexpression of different transcription factors [30,34,37,38,41]. These networks have been mechanistically analyzed at the systems level which led to the identification of regulators and destabilizers of different subtypes [30, 34, 38], and have contributed to our understanding of the underlying gene regulatory system. However, the structural properties of these networks were barely studied about a decade ago [42]. It has been shown in many studies that the structure of a network can be as important as its functional features and their analysis may help to identify key components associated with fundamental functional behaviors [43-45]. Specifically, *hubs* (Box 1) of the networks are shown to have key functional properties [46-51]. In this study, we topologically analyze the SCLC TF network (Figure 1) of [34, 38] that has been key in the identification of different SCLC subtypes. It comprises literature-based connections that are verified from ChEA, a database of ChIP-seq-derived interactions [52]. Overall, the network consists of 35 TFs connected through 239 activatory and inhibitory interactions (red and green arrows in Figure 1, respectively). Combinational ON–OFF states of the TFs in this network have been shown to drive cells toward different subtypes [34]. Here, one of our goals is to identify the hubs of the SCLC TF network, which are the special nodes that interconnect several key pathways and play an important role in collecting, processing, and distributing key signals throughout the signaling mechanism. We hypothesize that the hubs might be important for the overall network functioning and perhaps may help to identify specific TFs that regulate SCLC subtypes. Furthermore, although the earlier studies elucidate regulators of different SCLC subtypes, they lack mechanisms of subtype transitions whose understanding is critical to controlling disease progression. We also hypothesize that the pathways connecting the functionally distinct hubs may have roles in the subtype transitions.

**Figure 1.**
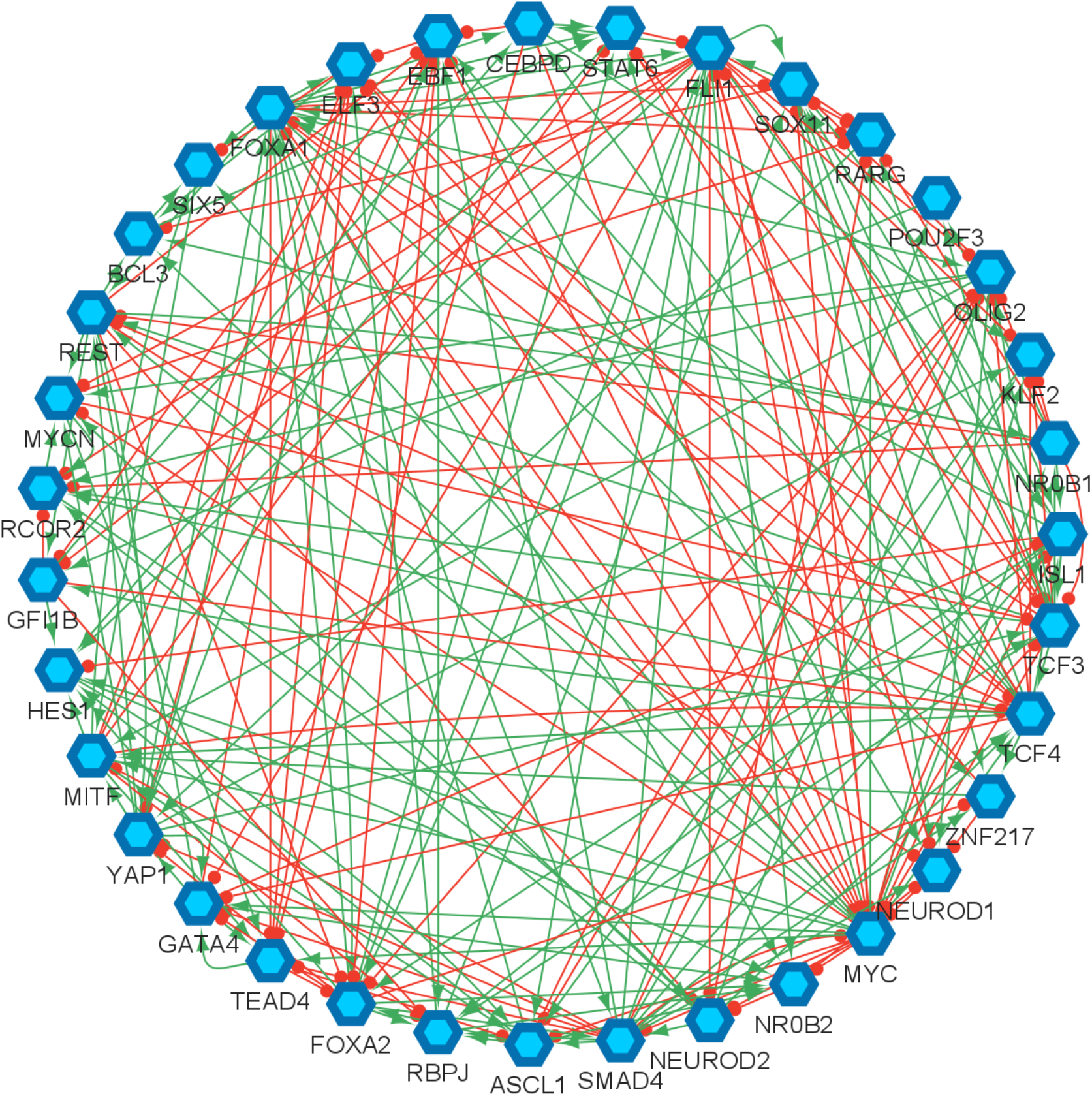
Small cell lung cancer transcription factor network reproduced from [34, 38]. The hexagonal nodes represent the individual transcription factors, the red edges represent the inhibitory interactions, and the green edges represent the activatory interactions.

To identify the hubs of the SCLC TF network, we implement a graph theory concept called Dense Spanning Tree (DST, see Box 1), which can be found by solving an optimization problem (Methods section A) [53-55]. We initially analyze a relatively unbiased network structure by considering the undirected and unweighted network. Later, we integrate previously-published RNA-seq data into our analysis, which is the probability of each interaction occurring [34, 38], assigned to each interaction as weights. To identify the hubs given the weighted network graph, we extend the DST concept into *Minimum Dense Spanning Tree* (MDST, see Box 1) concept for which the DST optimization problem is extended into a multi-objective optimization problem (Methods section B). Interestingly, all the found hubs are either regulators or destabilizers of the previously identified SCLC subtypes as elaborated in the Results section. Next, we test a pathway connecting the two functionally distinct hubs via simulations and observe a transition from the NON-NE to NE subtype. Furthermore, running and tracking several asynchronous NON-NE to NE transition simulations suggest additional TFs other than the hubs that may have a role in this transition.

The paper is organized as follows. First, we present the results of the DST and MDST analyses of the SCLC TF network in Results sections A and B. Then, we present the results of the asynchronous subtype transition simulations in Results section C. Next, we provide the mathematical details of DST and MDST analyses as well as the details of the transition simulations in Methods sections A, B, and C, respectively. In addition, we compare the DST and MDST analysis results in the Supplementary Material. Finally, we conclude the paper with some concluding remarks.

**Box 1: Brief Definitions**

- *Graph* is a collection of objects (points) linked together based on some pairwise relations. Figure B1-1 is an example of a graph (*G*) with the vertex set *V* = {a, b, c, d, e}. Some random weights are assigned to the edges for exemplary purposes.
- *Tree* is an acyclic graph, i.e., a graph that do not contain any cycles (loops). Figure B1-2 is an example of a tree.
- *Node (Vertex)* is an individual object (point) in a graph. “a” in Figure B1-1 is an example of nodes in the graphs.
- *Edge* is a link connecting two nodes in a graph. The link connecting “a” and “b” in Figure B1-1 is an example of edges.
- *Node Degree* is the number of edges connected to the node.

For more details on basic Graph Theory definitions, please see [56].

Given a graph *G* with a vertex set *V*:

- *Spanning Tree (ST)* is a subset of *G* that contains all the vertices in *V* with minimum number of edges [54]. They are not unique and known as the basis of the graph. Figure B1-2 is an example of ST. It contains all the vertices in *G* with minimum number of edges.
- *Minimum Spanning Tree (MST)* is a special spanning tree that minimizes the total weights assigned to the edges. Figure B1-3 is an example of MST. It is a ST and it minimizes the total edge weights.
- *Dense Spanning Tree (DST)*: is a special spanning tree that minimizes the total distances between the vertices [54]. Figure B1-4 is an example of DST. It does not care about the edge weights, but it minimizes the total distances between the nodes. Note that the distance between two nodes here is defined as the number of edges in the shortest path between the nodes, e.g., the distance between “a” and “e” in Figure B1-1 is two.
- *Minimum Dense Spanning Tree (MDST)*: is a special spanning tree of a weighted graph that minimizes the total distances between the vertices while minimizing the total weights assigned to the edges. Figure B1-5 is an example of MDST. It minimizes both total distances between the nodes and the total weights assigned to the edges.
- *Hub*: is a node (component) of a graph (network) that has the number of connections above average [57]. Node “b” in Figure B1-4 is an example for hubs, which has higher node degree and connects multiple nodes.

**Figure B1.**
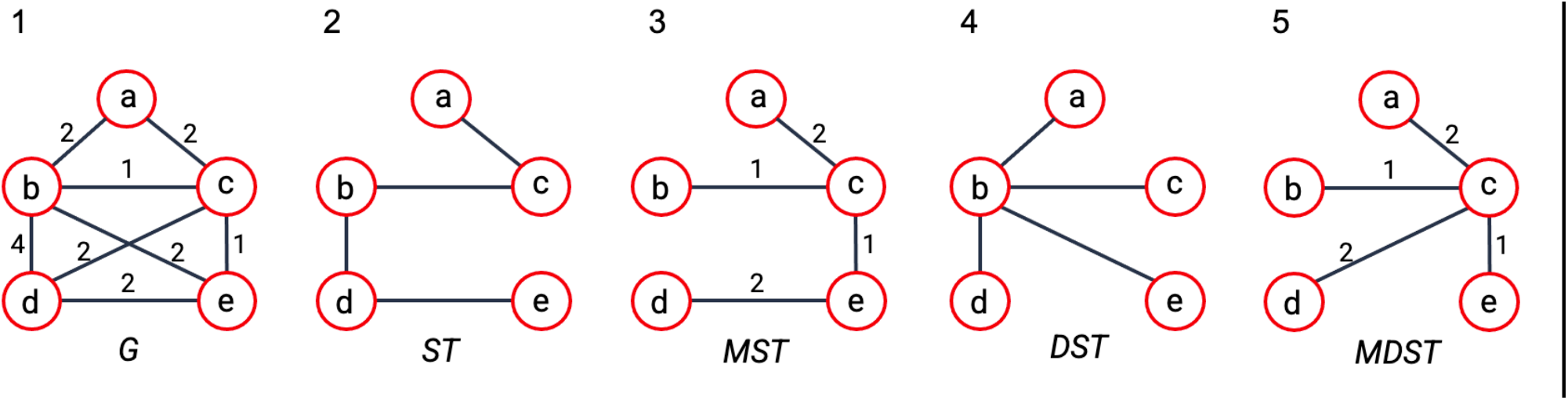
Examples for the introduced concepts. (1) An example of a weighted graph with random weights assigned for exemplary purposes. In a real network, the weight of an edge could be the likelihood (or strength) of the connection or other values such as mutual information, etc. (2) An example of a spanning tree. (3) An example of a minimum spanning tree. (4) An example of a dense spanning tree. (5) An example of a minimum dense spanning tree introduced in this paper (see Methods section B).

## RESULTS

In our analyses, given the SCLC TF network (Figure 1), we search for hubs of the network by finding the substructure DSTs (Box 1). The DST of a given network contains hubs that are known to be structurally important nodes interconnecting several pathways. Due to their high and strategic connectedness, they are very likely to have functional importance as well. This concept has many applications in different areas such as telecommunications networks, social networks, resource allocation, and biological networks [55].

In biological networks, the DSTs of the network are substructures that preserve the shortest pathways between the nodes (TFs) and hence they preserve the maximum influence among the individual components while highlighting a few nodes as the hubs. Since the identified hubs connect several pathways, they receive so many signals from their peripherals, process them, and distribute them to multiple other nodes. Therefore, in general, they have functional importance as well [46-51]. Also, depending on the size of the initial network, the identified DSTs may contain multiple hubs. Due to their individual importance, the pathways connecting the hubs might also be important as they are the pathways communicating complex signaling between the hubs. In this section, we show that the hubs of the SCLC TF network are relevant to the SCLC subtypes. Additionally, we test a pathway connecting two identified hubs via network simulations. All the results are elaborated in the following subsections.

### A. Structural analysis of the unbiased SCLC TF network identifies some of the known SCLC subtype regulators and destabilizers

We start our analysis by converting the SCLC TF network (Figure 1) into an undirected, unweighted network (see Methods section A). In this way, we just consider whether there is an interaction between two nodes or not without weighing their importance, which allows us to analyze a relatively unbiased network structure. Then, we search for the DSTs of the SCLC TF network following the approach of [55]. Upon solving the global optimization problem in Equation (1) (Methods section A), we observed 146,143 DSTs, all having the same optimum total distances between the TFs. Examples of the found DSTs are presented in Figure 2. In one of the DSTs, FLI1 and MITF are identified as the hubs (Figure 2A) while in the other DST, FLI1, ASCL1, and FOXA1 are identified as the hubs (Figure 2B). Since different DSTs may highlight different TFs as the hubs, we computed the average node degrees (Box 1) of the nodes among all the found 146,143 DSTs, which is collectively presented in Figure 3. As seen in the figure, FLI1 is a major hub with about 20 connections on average among all the found DSTs. In addition, MITF, ASCL1, NR0B1, and FOXA1 are the other hubs with relatively high average node degrees in some DSTs.

**Figure 2.**
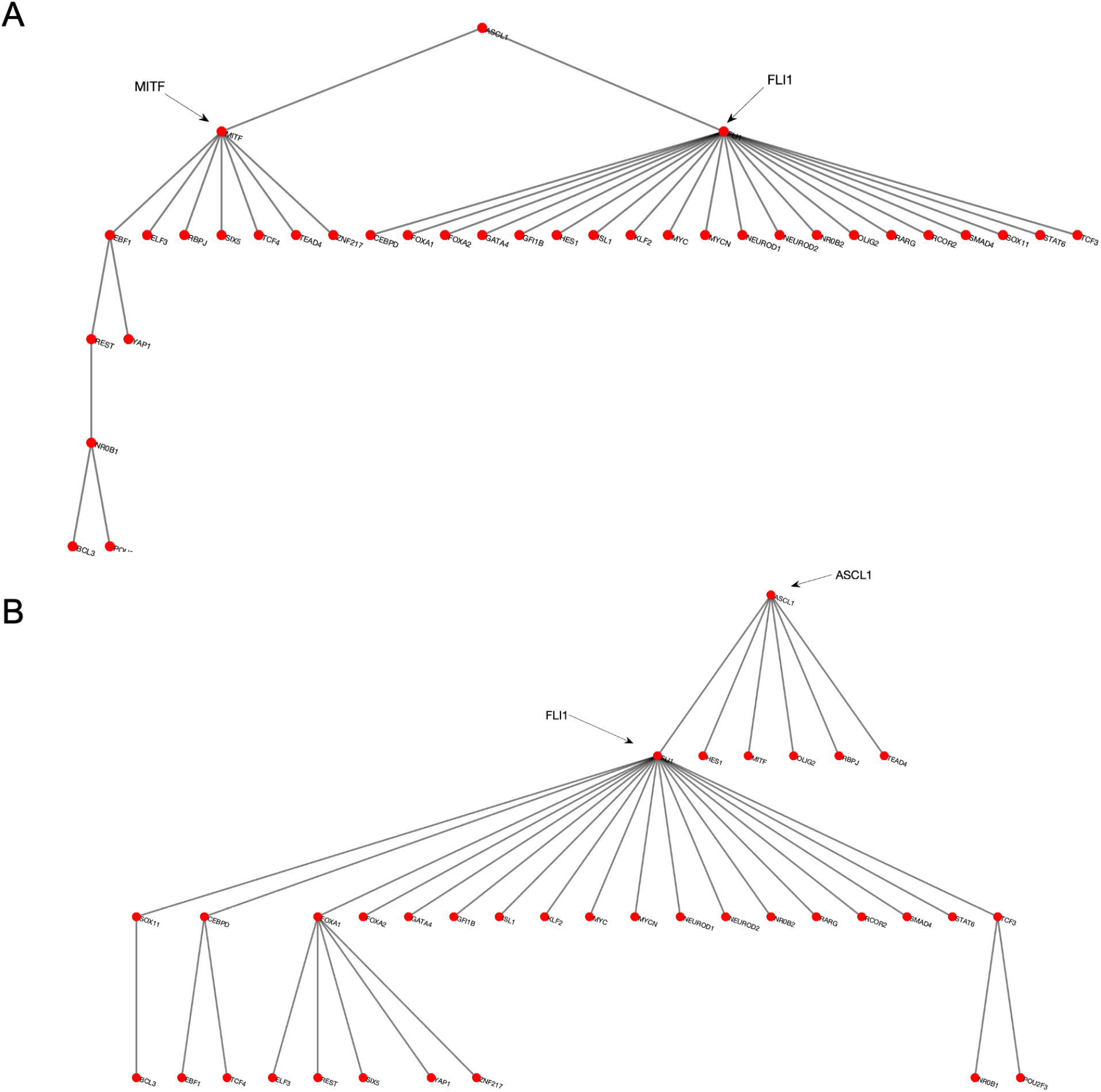
Examples of the found DSTs of SCLC TF network. (A) An example DST in which FLI1 and MITF are the two hubs. (B) An example of found DSTs in which FLI1, ASCL1, and FOXA1 are the three hubs.

**Figure 3.**
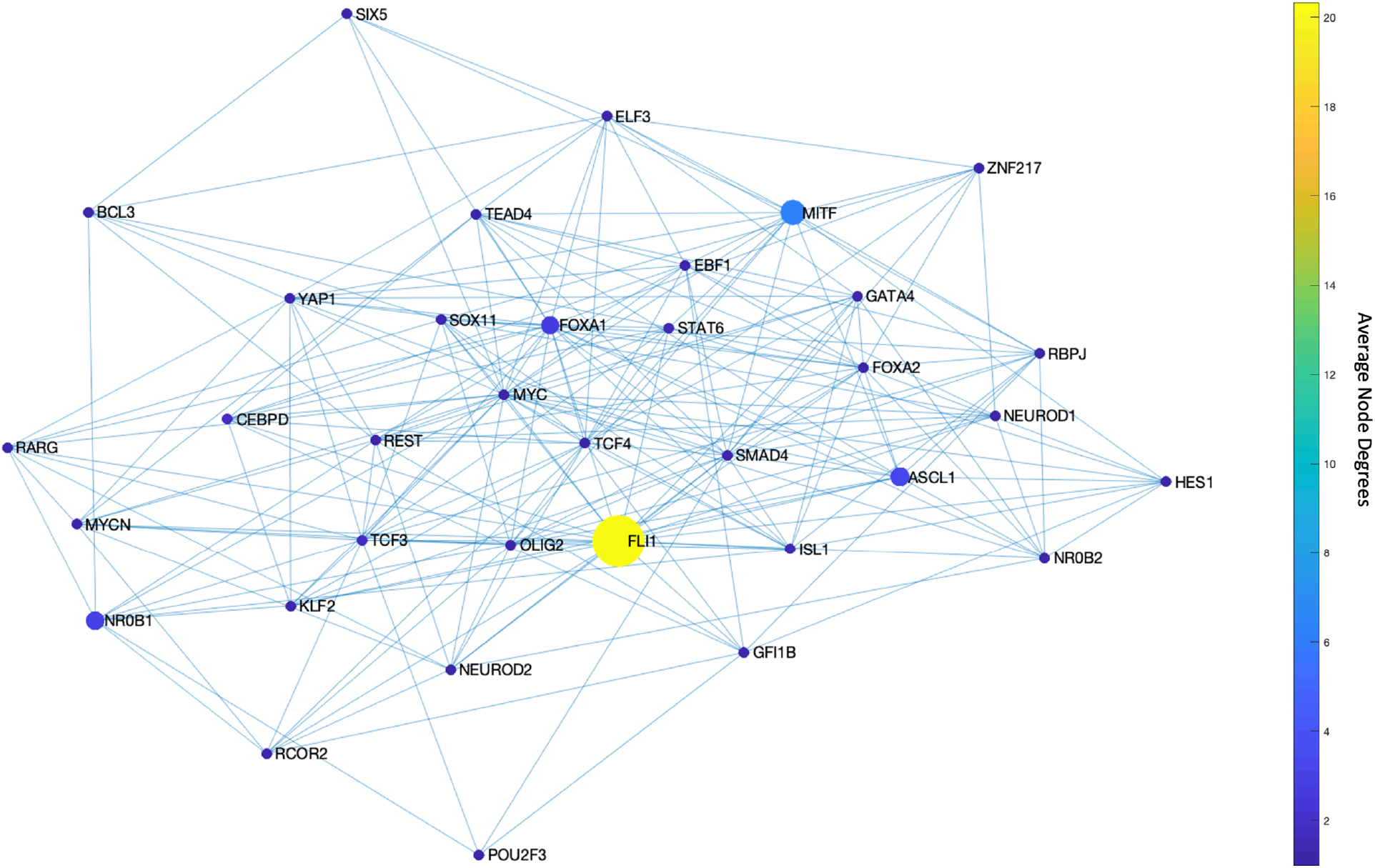
Average node degrees of each TF among the found DSTs. FLI1 is the major hub with about 20 connections on average in the found DSTs. The other hubs are MITF, ASCL1, NR0B1, and FOXA1 with relatively high connectedness on average.

The found major and side hubs are not only structurally important but also shown to have biological importance to the identified SCLC subtypes. For instance, FLI1 – the major hub in Figure 3 – is shown to be one of the regulators of the SCLC NE subtype [34,58,59]. Similarly, ASCL1, NR0B1, and FOXA1 are reported as one of the regulators of SCLC NE and NEv2 subtypes, and MITF is reported as one of the regulators of the SCLC NON-NE subtype [34], which shows the specificity of the hubs of SCLC TF network.

### B. Data-driven structural analysis of the SCLC TF network highlights MYC as a hub in addition to those previously identified as subtype regulators and destabilizers

Next, we repeat our hub search by integrating experimental data into the analysis. The data is the individual probabilities of each interaction between the TFs in the SCLC TF network (Figure 1), extracted from RNA-seq data [34]. The probabilities are integrated into the network structure as the weights that are assigned to the associated edges. Then, to identify the hubs of the weighted SCLC TF network, we extend the DST concept into MDST (Box 1) for which we solve an extended multi-objective optimization problem (Methods section B). Apart from DSTs, MDSTs allow us to highlight the hubs while preserving the maximum likelihood of the interactions.

Upon solving the optimization, we observed only 46 MDSTs which is drastically lower than the number of DSTs (146,143) found with the unbiased network structure. This means that this analysis guided by prior knowledge, i.e., the experimental data, can constrain the search space more efficiently. Once we compute the average node degrees among the found MDSTs, we observe that FLI1 still is the major hub (Figure 4). Similarly, ASCL1 and MITF are still identified as the hubs but this time with higher average node degrees compared to the unbiased network analysis (Figure 4). In other words, they become more major hubs, which coincides with their biological importance in SCLC as reported in the literature [30,31,34,38,40,60-62]. Interestingly, the data-driven structural analysis further reveals MYC as another hub (Figure 4), which does not appear in the unbiased network analysis (Figure 3). Recently, MYC was shown to be one of the key TFs for SCLC [32,63-65], which initiates Notch signaling to reprogram neuroendocrine fate from NE to NEv1 to NEv2 to NON-NE states [40]. Overall, our observations support that structurally important nodes are very like to be functionally significant as well. Therefore, such structural analyses could be an initial step in the analysis of complex intracellular networked processes because of their potential to pinpoint important network components, which would guide experimental target discovery.

**Figure 4.**
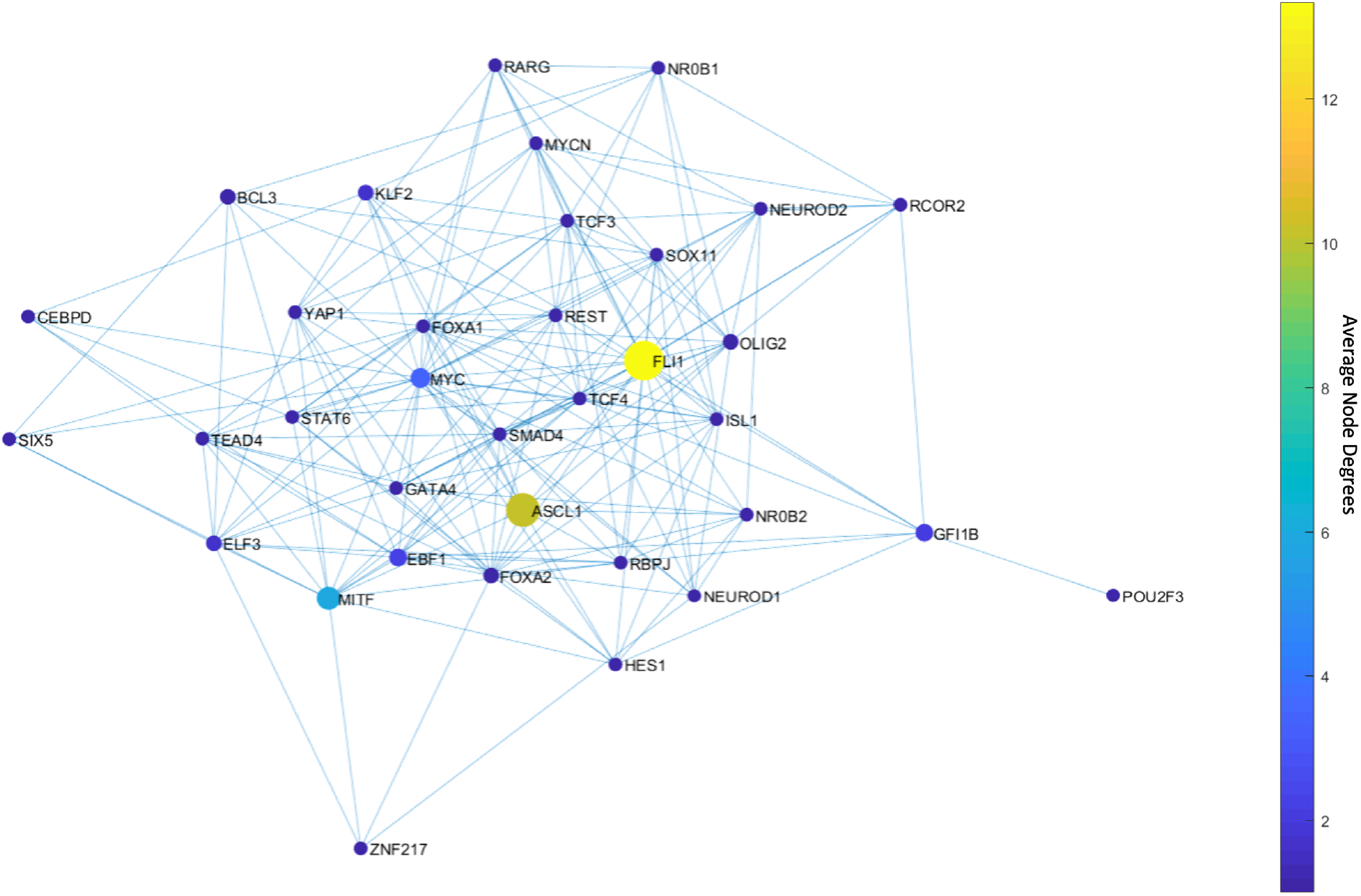
Average node degrees of each TF among the found MDSTs. FLI1 is the major hub with about 14 connections on average in the found MDSTs. The other hubs are ASCL1, MITF, and MYC with relatively high connectedness on average. Interestingly, MYC appears after integration of the data.

### C. The pathways connecting the SCLC TF network hubs may have a role in SCLC subtype transitions: NON-NE to NE transition occurs when FLI1 – ASCL1 – MITF pathway is active

SCLC TF network contains multiple hubs with varying average node degrees. These hubs are shown to have distinct functional features in terms of SCLC subtypes, as elaborated in the previous sections, which leads us to a question: Do the pathways connecting different hubs that are identified as regulators of different SCLC subtypes have any role in subtype transition? For instance, FLI1 and MITF are the two major hubs identified in both unbiased (Figure 3) and data-driven structural analyses (Figure 4). One of the pathways connecting these two hubs is through FLI1 – ASCL1 – MITF. FLI1 being a regulator of the SCLC NE subtype, MITF being a regulator of the NON-NE subtype, and ASCL1 being a destabilizer of the NON-NE subtype and regulator of the NE subtypes [34] suggest that this pathway has a potential role in NON-NE to NE subtype transition. One can also identify such structurally important pathways by checking the interactions remaining in the found DSTs and MDSTs with high probability, as exemplified in Supplementary Material.

To test the possible role of this pathway in the NON-NE to NE subtype transition, here we simulate the SCLC TF network using a tool called BooleaBayes [34] that automatically infers gene regulatory mechanisms, based on Boolean logic models, and links inputs and output states tailored to -omics datasets such as those from RNA-seq data. Upon setting the network’s initial state to NON-NE subtype based on previously identified combinational ON-OFF states of the TFs [34], keeping the FLI1 – ASCL1 – MITF pathway active, and running asynchronous network simulation (i.e., one TF is randomly picked and updated at each iteration) using the extracted logic rules (Methods section C), we observe a transition from NON-NE to NE subtype (Figure 5).

**Figure 5.**
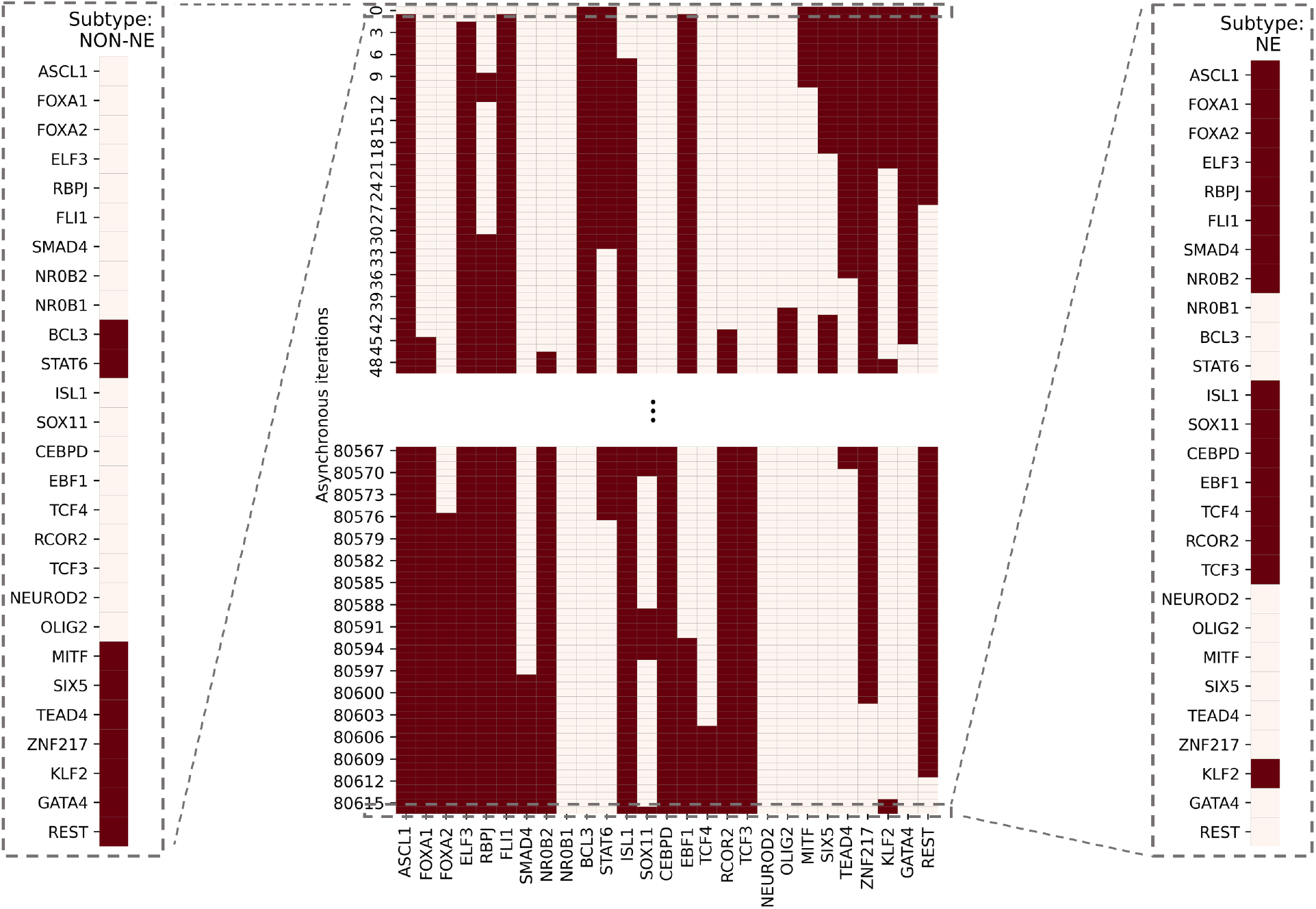
SCLC subtype transition from NON-NE to NE subtype. The network was initially set to NON-NE subtype. After running several asynchronous iterations by keeping the FLI1 – ASCL1 – MITF pathway active, the system converges to Ne subtype. This pathway was identified based on the hubs observed from both unbiased and data-driven network structure analyses. The details of network simulation are provided in Methods Section C. The red color means TF is ON and cream color means TF is OFF.

### Dynamic analysis of asynchronous NON-NE to NE subtype transition simulations

Although the NON-NE to NE subtype transition was observed by keeping the FLI1 – ASCL1 – MITF pathway active, there are possibly other TFs and dominant pathways that contribute to the transition. Identifying those TFs and dominant pathways may reveal how the system mechanistically executes such transitions and allow us to identify potential other TFs playing a role in the transition. Therefore, as the next step, we run 700 asynchronous NON-NE to NE subtype transition simulations and keep track of all the iterations. Then, we compute the Longest Common Sequence (LCS) based distance (Methods section D) between the target SCLC Boolean

NE state and the instantaneous network state at each iteration (Methods section C). As seen in Figure 6, throughout the NON-NE to NE transition, the network state dynamically alternates between NON-NE and NE subtypes through many distance-increasing and -decreasing patterns until it finally converges to the NE state. This means that some reaction patterns drive the cells toward the NE subtype (distance-decreasing patterns in Figure 7) whereas some other reaction patterns drive the cells toward the NON-NE subtype (distance-increasing patterns in Figure 7).

**Figure 6.**
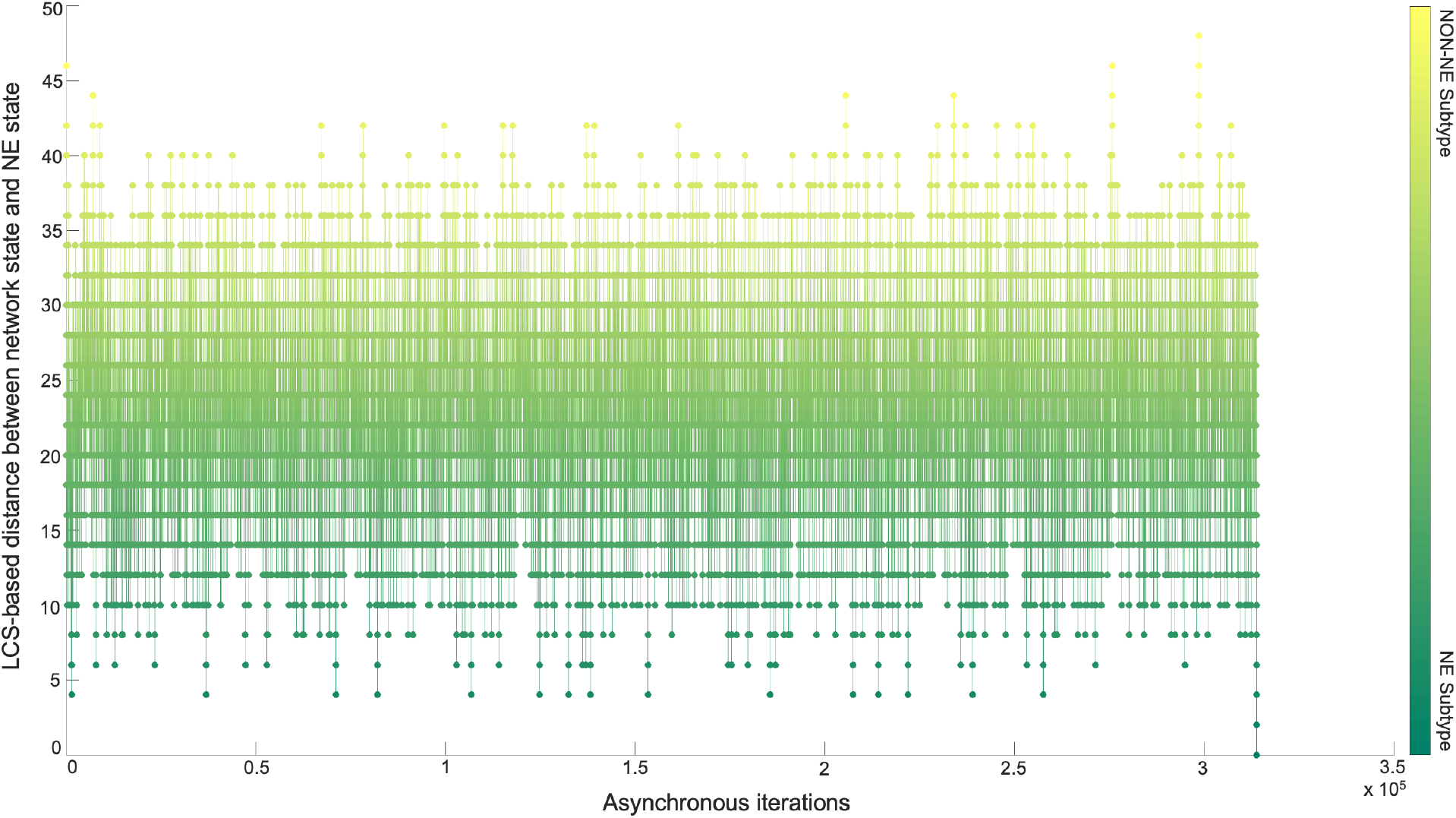
Longest Common Sequence-based distance between NE subtype and the instantaneous network state versus asynchronous iterations. Starting from NON-NE state, the system converges to and diverges from NE state multiple times throughout the iterations until finally it fully converges.

**Figure 7.**
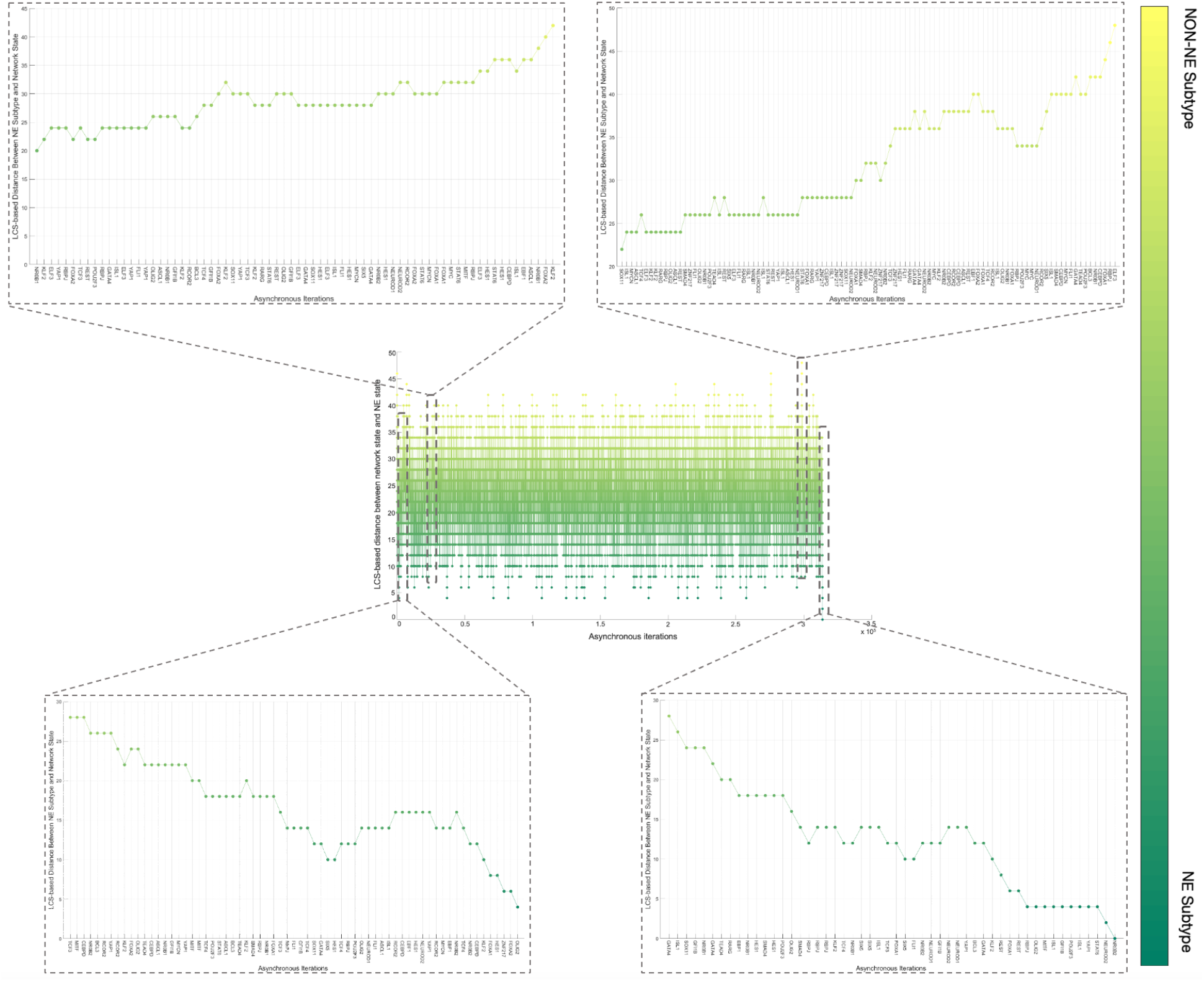
Examples of increase and decrease distance patterns between the network instantaneous state and SCLC NE subtype.

Overall, the 700 asynchronous NON-NE to NE subtype transition simulations, in which transition occurs in the order of 10^5^ asynchronous iterations, contain about 7×10^5^ distance increasing and 5×10^5^ distance decreasing patterns. To see which TF appears most in the distance-increasing and -decreasing patterns, we compute their frequencies (Figure 8). Interestingly, four TFs that are ASCL1, FLI1, NR0B1, and CEBPD, appear more than the other TFs in the distance-decreasing patterns (Figure 8A) whereas the same four TFs appear less than the others in the distance-increasing patterns (Figure 8B). This means that in addition to the ASCL1 and FLI1 which are part of the pathway identified NON-NE to NE transition pathway, NR0B1 and CEBPD may have a regulatory involvement in this transition as well. Moreover, throughout all the asynchronous iterations among 700 NON-NE to NE transitions, we compute the number of iterations for each TFs, on which an update of the TF causes an increase in the distance between the network’s instantaneous state and NE subtype. As seen in Figure 9A, in addition to ASCL1 and FLI1 which never drives the cells toward the NON-NE subtype, NR0B1 and CEBPD are the two TFs that have a lower effect on the increase in the distance between the network state and the NE subtype compared to the others, which further supports their possible regulatory involvement in NON-NE to NE subtype transition. Furthermore, we compute the probability of TFs being ON at the network state during the initiation of distance decrease patterns (Figure 9B). With about 0.2 probability of being ON, NR0B1 seems to drive the cells toward the NE subtype by mostly being OFF whereas the activity status of CEBPD seems not very important as its probability of being ON is very close to 0.5. Additionally, Figure 9B suggests that whenever ISL1 and FOXA2 appear in the distance-decreasing patterns which is very likely as seen in Figure 8A, they are mostly ON with relatively high probabilities which implies that they may have a role in the NON-NE to NE transition.

**Figure 8.**
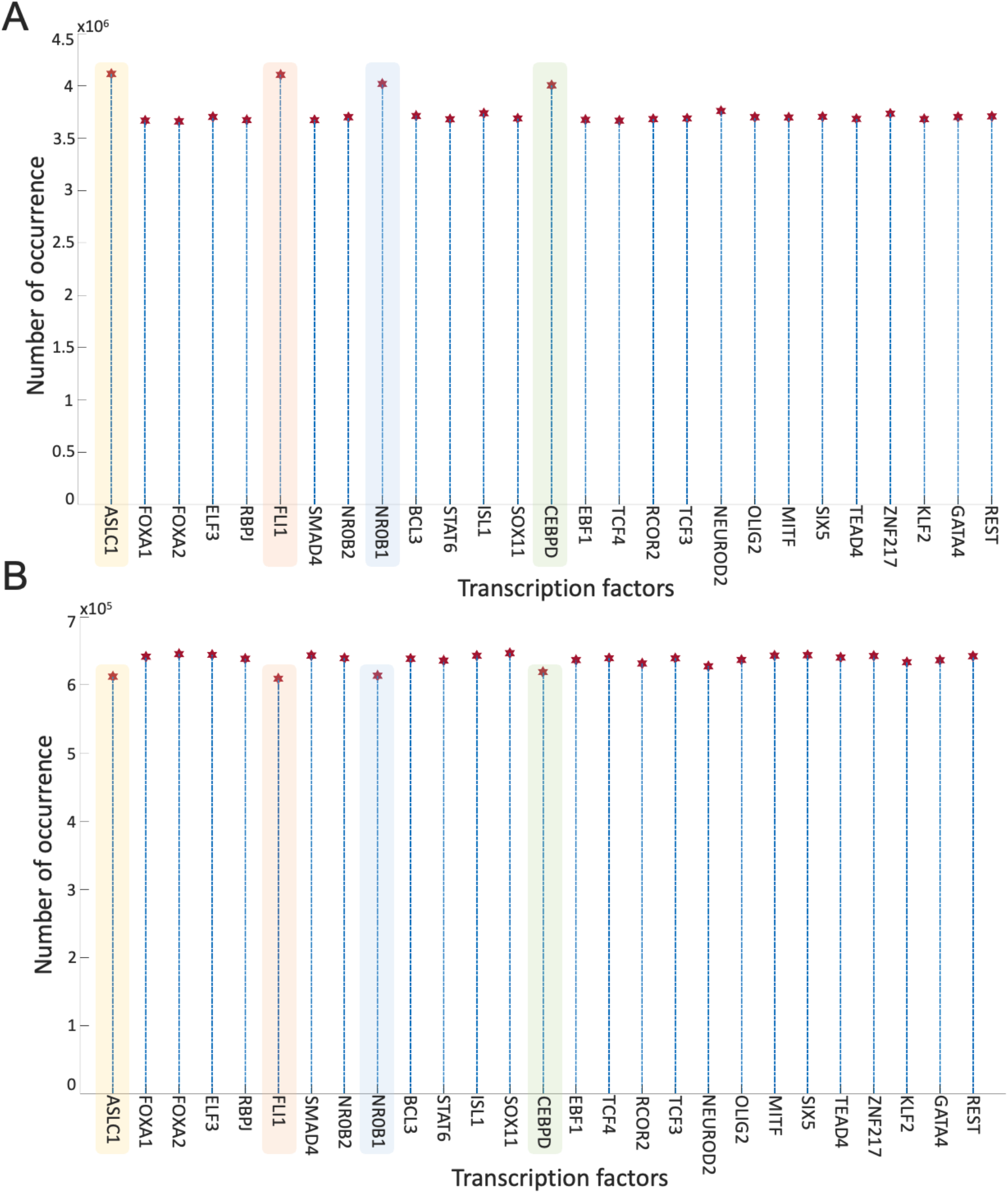
Frequencies of TFs in the distance decreasing and increasing patterns. (A) Appearance of TFs in the distance decreasing patterns. (B) Appearance of TFs in the distance increasing patterns.

**Figure 9.**
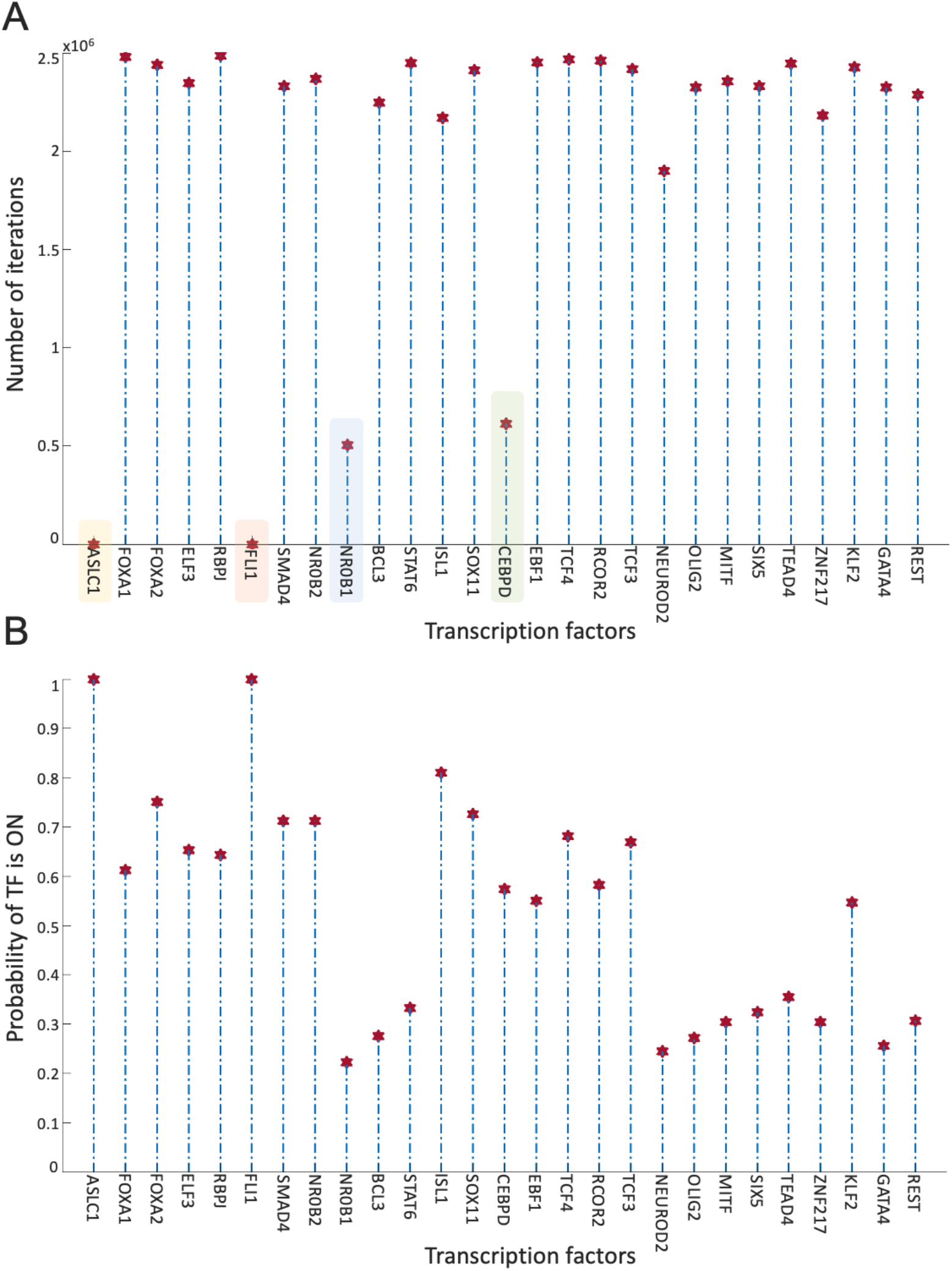
Effect of TFs in distance increase and decrease between network state and NE subtype. (A) Number of iterations on which update of TF cause an increase in the distance between network state and NE subtype. (B) Probability of TF is ON in the network state initiating distance decreasing patterns.

Overall, the presented results suggest that structural analysis of the biological networks may guide the identification of functionally important molecules. More specifically, the concepts of DST and here extended to MDST by integrating data can identify hubs of the networks which can be potential targets in the experiments due to their involvement in complex biological processes. Focusing on the SCLC TF network that is being analyzed in this work, all the identified hubs in both unbiased and data-driven analysis show biological importance in terms of SCLC subtype regulation and destabilization as supported by the literature. Moreover, integrating data into the structural analysis highlights MYC as another hub whose importance in SCLC subtypes has recently been discovered [32,63-65]. This observation further supports those previously reported results. Furthermore, the ability to identify multiple hubs that have distinct functional roles in SCLC subtypes lets us scrutinize the pathways connecting the hubs. Upon asynchronously simulating the network by keeping the pathway connecting FLI1 and MITF – the two major hubs – active, we observed a transition from NON-NE to NE subtype. In addition, analysis of 700 asynchronous NON-NE to NE transition simulations suggests other TFs that may play a role in this transition. As a result, starting from a pure network structure, its analysis leads us to understand the underlying mechanism of a complex biological system, which is noteworthy.

## METHODS

### A. Dense Spanning Trees of the unbiased SCLC TF network

Given the SCLC TF network (Figure 1), to analyze its structure and identify the hubs (Box 1) that are potentially fundamental in terms of their roles in complex biological processes, we search for the substructures called dense spanning trees (DSTs, Box 1). Suppose *G* is a graph that represents the SCLC TF network, *V(G)* is the set of nodes that represent the TFs in the network and *E(G)* is the set of edges that represents the interactions between the TFs in the network. Then, the DST of *G* is a substructure that minimizes the total distances between the TFs and contains all the TFs in *V(G)* with a minimum number of interactions while highlighting some nodes with high connectedness, i.e., the hubs. In other words, the DSTs are the subnetworks of the SCLC TF network that comprises the hubs and the shortest pathways from the hubs to all other TFs preserving the maximum biological influence.

To identify the hubs of the SCLC TF network, we start with a relatively unbiased network structure by removing all the edge directions, I.e., the information on activatory and inhibitory interactions, and not using any data on strength of the connections (Supplementary Figure 1). Then, the DSTs of the network are observed by solving the following optimization [55]:

For the graph *G* with vertex set *V*(*G*) = {*v*_1_, *v*_2_, …, *v*_*N*_} where *N* = |*V*|, and edge set *E*(*G*) = {*e*_1_, *e*_2_, …, *e*_*M*_} where *M* = |*E*|,

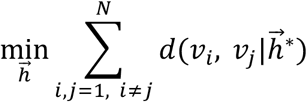

subject to

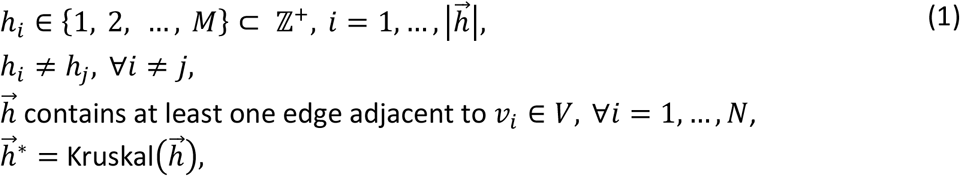

in which 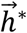denotes the minimum spanning tree obtained from 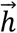 that is a subset of *E(G)*, and *d*(*v*_*i*_, *v*_*j*_) is the distance between nodes *v*_(_ and *v*_)_ defined as the total number of edges in the shortest pathway between *v*_(_ and *v*_)_. The main idea here is to find the optimal subset(s) of edges *E(G)* from which the constructed DST has the optimal objective value which is the total distances between the individual nodes. For more mathematical details and possible applications of this approach, we refer the reader to [54,55]. Upon solving the optimization problem (1) via Genetic Algorithm (GA), which is a metaheuristic optimization method that attempts to find the global optimum or at least its good approximation [66], we observed 146,143 DSTs with the same objective value.

### B. Integrating data into the structural analysis: Minimum Dense Spanning Trees

As the next step, we blend this pure structural analysis with some data that is the probability of the existence of the interactions, i.e., the strength of the connections estimated from RNA-seq data [34]. The probabilities are integrated into the network structure as the weights that are assigned to the associated edges. Then, to identify the hubs of the weighted SCLC TF network, here we reformulate the optimization problem constructed to find DSTs in Equation (1) as a multi-objective optimization problem given in Equation (2) and call the resulting optimal trees as the minimum dense spanning trees (MDSTs, Box 1). MDSTs add another information layer to the found trees by preserving the maximum likelihood of the interactions in addition to the minimum total distances between the nodes while highlighting the hubs of the network. More precisely, MDSTs of the SCLC TF network are the subnetworks that preserve the most probable interactions as well as the maximum biological influence between the TFs via the shortest pathways through the hubs. Note that one can assign different weights to the interactions by different means such as the mutual information between the TFs extracted from experimental data. In this case, the MDSTs will be the substructures that preserve the highest mutual information in addition to the shortest pathways through the hubs.

To find the MDSTs of the SCLC TF network, we extend Equation (1) as follows: Suppose for each interaction *i*, we are given a probability *p*_*i*_, that is probability of the existence of the *ith* interaction. Then, for the graph *G* with vertex set *V*(*G*) = {*v*_1_, *v*_2_, …, *v*_*N*_} where *N* = |*V*|, and edge set *E*(*G*) = {*e*_1_, *e*_2_, …, *e*_*M*_} where *M* = |*E*| with associated weights *w*_(_, *i* = 1, …, *M*:

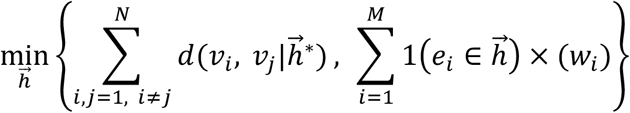

subject to

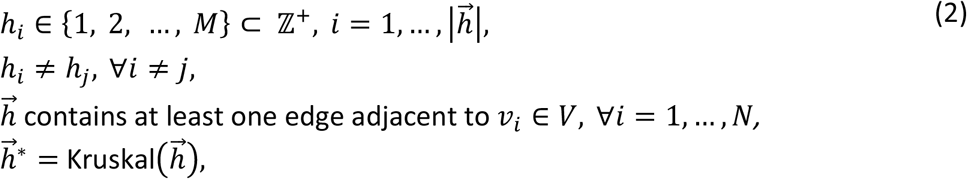

in which weight 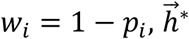 denotes the minimum spanning tree obtained from 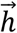that is a subset of *E(G)*, and *d(v*_*i*_, *v*_*j*_) is the distance between nodes *v*_*i*_ and *v*_*j*_, and 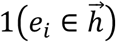 results in 1 if the edge *e*_*i*_ is in 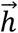. Here, the first objective function is the minimization of the total sum of distances between the nodes whereas the second objective function is the minimization of the sum of weights assigned to each edge, which is the same as the maximization of the sum of probabilities of each selected interaction exists based on the definition of weights. Once we solved the multi-objective optimization problem (2) by GA, we observed 46 MDSTs all having the same objective value, which shows the effect of prior knowledge on narrowing down the search space.

### C. SCLC TF network subtype transition simulations

To see how important the pathways connecting the hubs having distinct functional features are, we simulate the SCLC TF network using a tool called BooleaBayes [34]. BooleaBayes is a Boolean rule-fitting algorithm that infers local regulatory mechanisms near stable cell subtypes from gene expression data. The approach has previously been applied to the SCLC TF network (Figure 1) to identify and rank master regulators and master destabilizers of SCLC subtypes assuming binary, i.e., ON and OFF, activity states of each transcription factor (Supplementary Figure 2). Further details of BooleaBayes and how it infers the logic rules can be found in [34].

Using the Boolean rules extracted via BooleaBayes, we test the role of FLI1 – ASCL1 – MITF pathway, in which FLI1 and MITF are the two major hubs found by both DST and MDST approaches, in NON-NE to NE subtype transition. This is hypothesized due to FLI1 being a regulator of the SCLC NE subtype, MITF being a regulator of the NON-NE subtype, and ASCL1 being a destabilizer of the NON-NE subtype and regulator of the NE subtype [34]. First, we set the initial state of the network to the NON-NE subtype using the logic TF states in Supplementary Figure 2. Then, we simulate the network using a general asynchronous update scheme with the inferred Boolean rules and keeping the FLI1 – ASCL1 – MITF pathway active by setting ASCL1 and FLI1 always “ON”. After several asynchronous iterations (usually in the order of 10^5^), in which a random TF is picked at each iteration and updated based on the extracted probabilistic Boolean rules, the network converged to one of the NE subtype Boolean states (Supplementary Figure 2). Note that due to the nature of the asynchronous update scheme, the convergence of the system to the NE subtype may occur in a different number of iterations and update patterns at each run of the transition simulations.

### D. Distance measure between instantaneous network state and NE subtype

To track the network state and understand its dynamic behavior throughout NON-NE to NE transition, we compute the distance between the network’s instantaneous state at each iteration and the target NE subtype. The distance metric we chose is Longest Common Sequence (LCS) metric [67] due to its sensitivity to order differences by assigning a larger distance value to the difference between the network state and target state. Given two vectors *v*_1_ and *v*_2_ of length *m*, that in our case represent the network state and the target state, respectively, the LCS-based distance *d*_LCS_ is defined as follows:

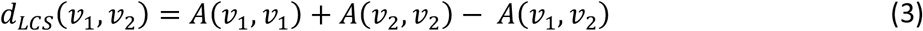

where *A*(*v*_1_, *v*_2_) is to the number of elements in *v*_1_ that uniquely matches the elements of *v*_2_ in the same order (not necessarily contiguous). Note that one can use other distance metrics such as Hamming distance to perform the same analysis.

Computing LCS-based distance between the instantaneous network state and NE subtype throughout the asynchronous transition simulations shows us how the network converges and diverges from the NE subtype starting from the NON-NE subtype. Furthermore, this allows us to identify some patterns causing increase and decrease between the two network states; and hence, allows us to identify other TFs that may contribute to this transition.

## DISCUSSION

Small Cell Lung Cancer (SCLC) is an aggressive disease with its mixtures of transcriptional subtypes such as *neuroendocrine* (NE) and *non-neuroendocrine* (NON-NE), later being more treatment-resistant, regulated by the expression of different transcription factors (TFs). In addition to the heterogeneity in cancerous cell types, transitions between the subtypes make the disease even harder to treat. To date, SCLC TF networks have been broadly studied via systems approaches to reveal regulators and destabilizers of different subtypes. Yet, the studies lack mechanisms of subtype transitions, whose understanding is critical to control disease progression and perhaps develop ways for permanent cure. In this work, we hypothesize that analysis of the SCLC TF network structure (Figure 1), which is barely investigated to our best knowledge, can provide clues on distinct subtype drivers, and further reveal pathways controlling subtype transitions. To test this hypothesis, here we use graph theory concepts called Dense Spanning Trees and its extended version called Minimum Dense Spanning Trees (DSTs and MDSTs, see Box 1 and Methods sections A and B). DSTs and MDSTs are special subnetworks of the initial TF network that feature strategical nodes called hubs and the pathways connecting the hubs. Hubs are critical nodes due to interconnecting several key pathways and collecting, processing, and distributing key signals throughout the signaling mechanism. Moreover, the pathways connecting the hubs are also important as they are potential probes for controlling complex signaling across hubs. Therefore, given two hubs regulating different SCLC subtypes, we hypothesize that the pathways connecting these hubs could be targets to control subtype transitions.

First, with DSTs, we analyze a relatively unbiased network structure by removing all the edge directions, i.e., the information on activatory and inhibitory interactions, and not using any data on strength of the connections (Figure 3). Next, we integrate data into this pure structural analysis, assigned to each edge as weights that are the probability of the existence of the interactions, i.e., the strength of the connections estimated from RNA-seq data [34]. Then, we extend the DST into MDST (Methods section B) to identify the hubs of the weighted network structure (Figure 4). Interestingly, all the hubs such as ASCL1, FLI1, and MITF identified in both unbiased and data-driven structural analyses are either regulators or destabilizers of different SCLC subtypes as reported in the literature, which confirms our hypothesis on the importance of hubs. Additionally, the structural analysis driven by the data highlights MYC as another hub in addition to those identified in unbiased analysis (Figure 4), which supports its importance in SCLC subtypes as shown in recent studies [32,63-65]. To test the roles of pathways connecting functionally distinct hubs, we asynchronously simulate the SCLC TF network using a Boolean modeling framework extracted by a tool called BooleaBayes [34] (Methods section C). As a result of several asynchronous iterations and keeping the pathway connecting FLI1 and MITF – the two major hubs in both unbiased and data-driven analyses – active, we observe a transition from NON-NE to NE subtype (Figure 5), confirming our hypothesis on the importance of hub-connecting pathways. Furthermore, after analyzing increasing and decreasing patterns in distance between the network state and NE subtype (Figure 6 and Figure 7) in 700 asynchronous NON-NE to NE transition simulations, we conclude that the TFs NR0B1 and CEBPD may also play a role in this transition in addition to FLI1 and ASCL1 (Figure 8 and Figure 9).

Note that, one can integrate different data into this analysis, assigned as the weights to the edges. For instance, instead of assigning probabilities of interactions, the mutual information between the pair of nodes can be used. In this case, resulting MDSTs would contain the hubs while preserving the highest mutual information and the maximum influence within the nodes. Similarly, one can assign the weights manually guided by prior knowledge to keep the preferred interactions in the resulting substructures. Also, one can apply the tools presented here for any network type such as protein-protein interactions networks (PPINs), gene regulatory networks (GRNs), cell signaling networks, and metabolic networks. In addition, they can be applied to any network structures such as directed or undirected and weighted or unweighted. We would like to note that although preserving the directedness of interactions would integrate more information into the structural analysis, it would also require adding new constraints to the optimization problems (1) and (2), which may become harder to solve due to increased complexity, leaving room for potential improvement to the found DSTs and MDSTs for the SCLC network.

There are different ways to define and identify the hubs for a given network than ours. One can define a node that has the most connections (highest node degree) or a node that has the most connections that make it central in the network as the hub. However, we believe they are not very well suited for biological applications as they are purely structural concepts and don’t concern about the closeness, i.e., the influence of the nodes with each other. Moreover, such hubs are expected to occur only in scale-free networks, i.e., the networks whose degree distribution follows power law [57]. On the other hand, the concept of DSTs and MDSTs can identify hubs for any given network because, in DSTs and MDSTs, hubs are defined as the central nodes that minimize the total distance between every node, and such substructures can be found for any random network. Additionally, there are other ways to find DSTs of a given network such as the edge-swap heuristic algorithms presented in [53, 54]. However, we have previously shown that optimization-based approaches outperform such edge-swap heuristic algorithms [55] both in accuracy and computational complexity changing by the network size. Lastly, here, to identify the DST and MDSTs, we solve the optimization problems (1) and (2) using genetic algorithm (GA), which is a metaheuristic optimization method that attempts to find a globally optimal solution, but it does not guarantee a global solution because it does not guarantee exploration of all the search space and the solution quality and optimality depend on several parameters that need to be properly selected by the user, including population size, rate of mutation and crossover, etc. [66]. However, GA is well suited for problems that are discrete and combinatorial in nature by providing at least a good approximation of the global solution. Nevertheless, one can solve these optimization problems via other algorithms such as particle swarm optimization.

Overall, the presented results have shown that the hubs of the SCLC TF network identified via DSTs and MDSTs are either regulators or destabilizers of different SCLC subtypes. This implies that structural analyses of the networks can be advantageous as the initial step as their results can be used as guidance to generate hypotheses to be tested in experiments. Moreover, the pathways connecting the functionally distinct hubs may have major roles in SCLC subtype transitions as shown by the simulations, which may allow the control of such transitions and help develop better treatment strategies by driving the cancerous cells toward more sensitive states. Furthermore, targeting those pathways in the experiments may lead to the identification of other dominant components in such transitions and hence help to understand the underlying mechanism of this complex signaling process. As a result, pure as well as data-driven structural analyses of the networked processes could be a plausible first step and may result in potentially important biological observations in complex systems as well as help generate hypotheses to be tested.

## Acknowledgements

The authors would like to thank Vito Quaranta, Sarah Maddox Groves, and Lopez Lab members at Vanderbilt University for insightful conversations and critical feedback on this work. This work was supported by the following funding sources: CFL was supported by the National Science Foundation (NSF) [MCB 1411482] and NSF CAREER Award [MCB 1942255]; and the National Institutes of Health (NIH) [U54-CA217450 and U01-CA215845].

