## Supplementary document for "Data-driven structural analysis of Small Cell Lung Cancer transcription factor network suggests potential subtype regulators and transition pathways"

**DST versus MDST: Comparing interactions remaining on the found DSTs and MDSTs**

The dense spanning trees (DSTs) of the SCLC TF network are the substructures that emphasize some TFs as the hubs while preserving minimum total distances between the TFs and hence the maximum influence on each other. On the other hand, MDSTs of the weighted SCLC TF network are the substructures that still emphasize some TFs as the hubs and preserve the maximum influence between the TFs while minimizing the total weights assigned to the edges, that is for each edge $e_i$, the weight $w_i = 1 - P(e_i \text{ exists})$. Once we solved the associated optimization problems (Equations (1) and (2) in the main text), we observed 146,143 DSTs and 46 MDSTs all having the same objective values for their associated objective functions. Looking at the average node degrees among all the found DSTs and MDSTs, we have seen that most of the found hubs overlap between both analyses.

Here, we compare the interactions remaining in the DSTs and MDSTs. To do so, we computed the probability of an interaction remaining in the found DSTs (Figure S3A) and MDSTs (Figure S3B). As seen in the figure, some edges always remain in the found DSTs and MDSTs. For instance, the interaction between FLI1 and MITF always remains in the found DSTs. Similarly, the interaction between MITF and EBF1 always remains in the found MDSTs. Upon comparing all the interactions that always remain in the found DSTs and MDSTs, i.e., $P(e_i \text{ remaining in DST and MDST}) = 1$, we have seen that the interactions ASCL1–FLI1, GATA4–FLI1, ISL1–FLI1, MYCN–FLI1, NEUROD1–FLI1, NEUROD2–FLI1, RARG–FLI1, RCOR2–FLI1, SOX11–FLI1, STAT6–FLI1, and TCF3–FLI1 are common. This means that to observe the minimum total distance and maximum influence between the TFs network, these interactions should be kept in the DSTs and MDSTs, which shows their structural importance.

Additionally, the interactions having a high probability of remaining in the found DSTs and MDSTs might help to identify the possible important pathways between the hubs. For example, the interaction between the ASCL1–FLI1 always remains in both DSTs and MDSTs. Also, the MITF–ASCL1 connection has a probability of 1 for DSTs and 0.8 for MDSTs, meaning that it is very highly likely to have this connection in both substructures. This means that it is highly probable that the pathway FLI1–ASCL1–MITF also exists in the found DSTs and MDSTs, in which FLI1 (regulator of NE subtype) and MITF (regulator of NON-NE subtype) are two major hubs. Therefore, one can target this pathway both in silico and in experiments to test their potential impact on SCLC subtypes and NON-NE to NE subtype transitions as done in the main text.

### Supplementary Figures

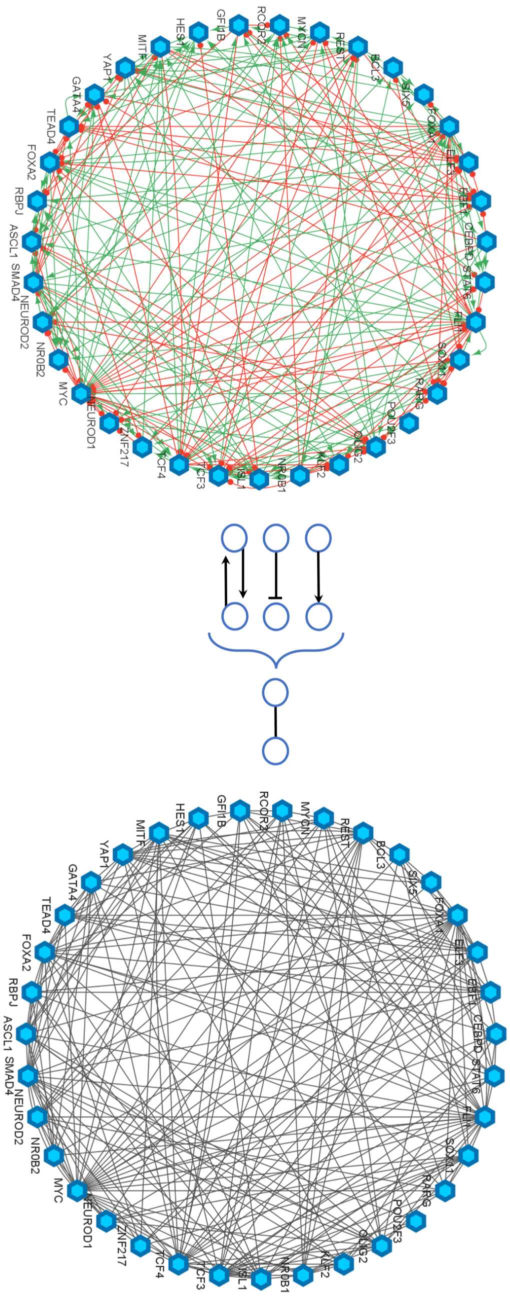


Figure S1. Converting directed SCLC TF network into undirected network to observe relatively unbiased network structure. Here, we only care whether there is an interaction between the two TFs and ignore the type of interaction, i.e., activation or inhibition.

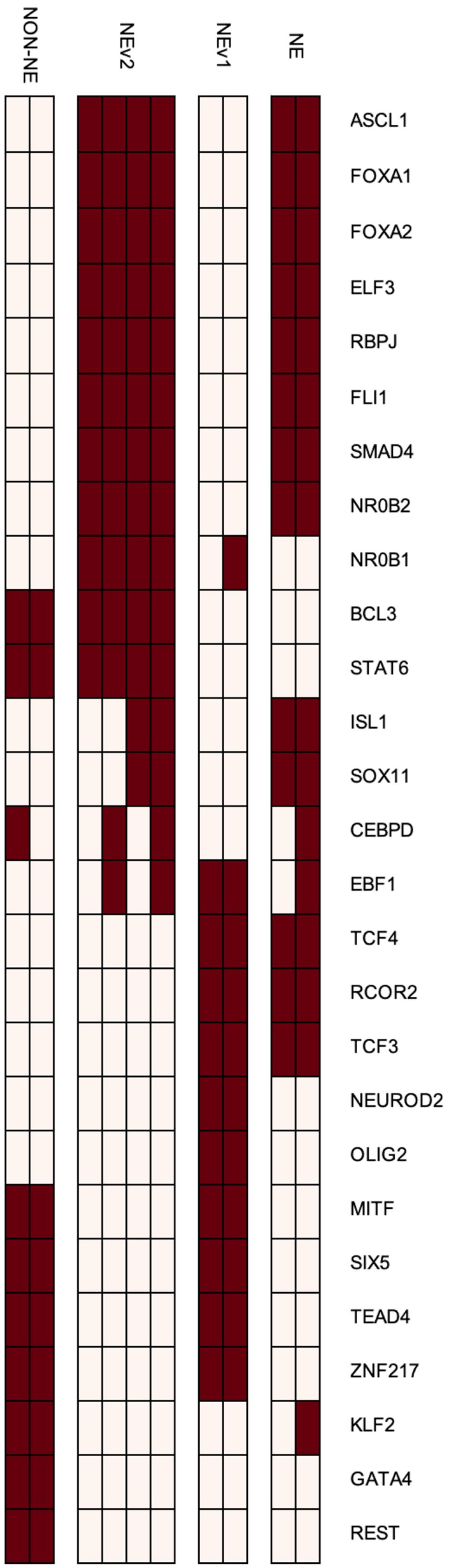


Figure S2. Boolean states of each TFs in different SCLC subtypes as identified by Wooten et al. [34].

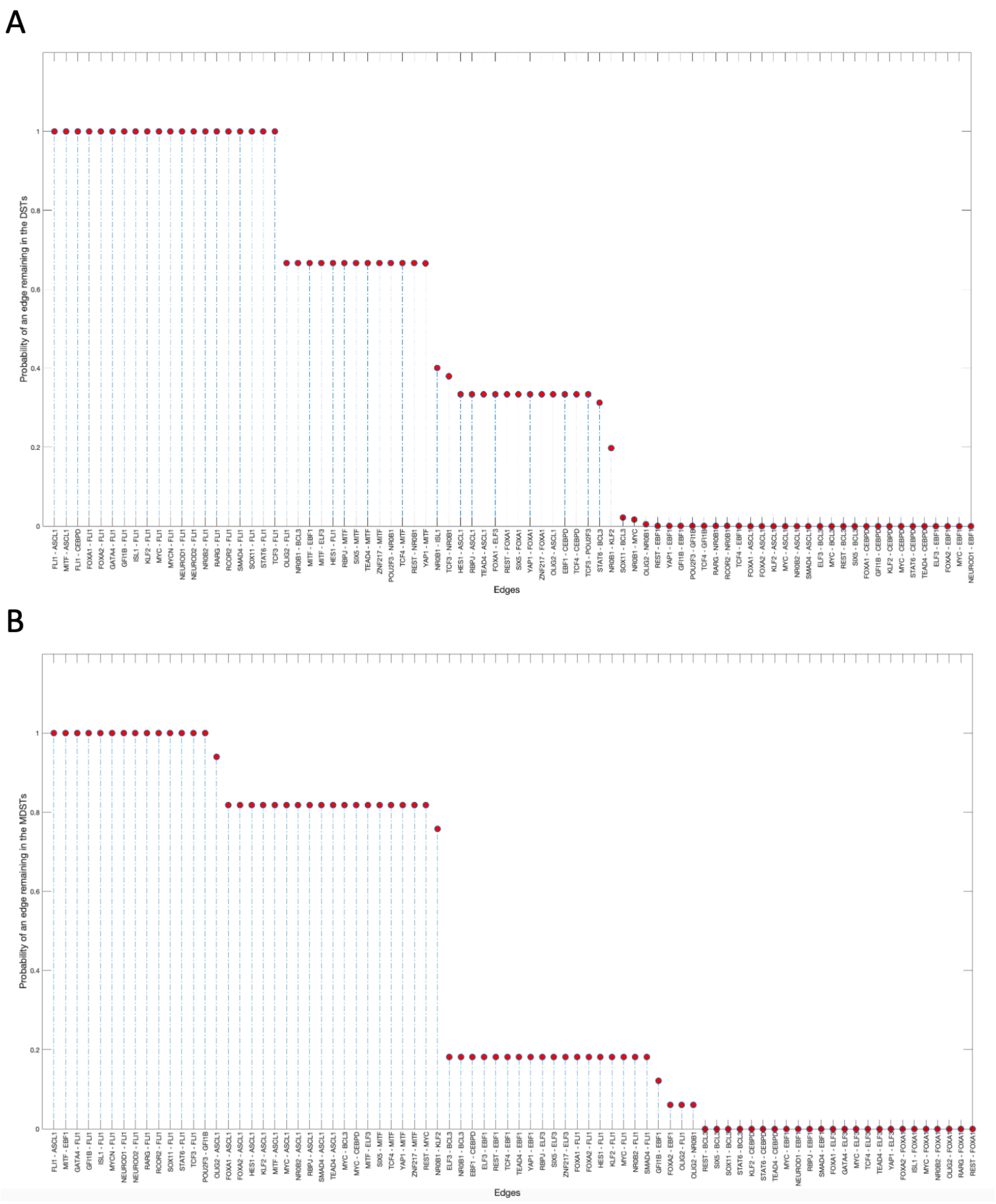


Figure S3. Probabilities of interactions remaining in the found DSTs and MDSTs.
